# Dynamic off-resonance correction improves functional image analysis in fMRI of awake behaving non-human primates

**DOI:** 10.1101/2023.10.18.562796

**Authors:** Mo Shahdloo, Nima Khalighinejad, Luke Priestley, Matthew Rushworth, Mark Chiew

## Abstract

Use of functional MRI in awake non-human primate (NHPs) has recently increased. Scanning animals while awake makes data collection possible in the absence of anaesthetic modulation and with an extended range of possible experimental designs. Robust awake NHP imaging however is challenging due to the strong artifacts caused by time-varying off-resonance changes introduced by the animal’s body motion. Recently, an image reconstruction technique has been proposed to estimate these off-resonance changes using the navigator data that is typically collected during fMRI scans to correct the data and compensate for the changes. In this study, we sought to thoroughly investigate the effect of this correction on the brain activation estimates using extended awake NHP data. Our results show significant improvements in image fidelity using our proposed correction strategy, as well as greatly enhanced and more reliable activation estimates in GLM analyses.

## Introduction

Neuroanatomical and functional parallels between humans and nonhuman primates (NHPs) have made NHPs useful models for understanding the human brain (Andersen et al., 2019; Gray and Barnes, 2019; Picaud et al., 2019; Roberts and Clarke, 2019; Rudebeck et al., 2019). However, robust functional magnetic resonance imaging (fMRI) of the NHPs at high spatiotemporal resolution is challenging due a variety of factors, including the anatomical differences in brain size and head shape, behavioural differences in the scanner, and the specialized receive coils needed to facilitate NHP data acquisition (Autio et al., 2021; Friedrich et al., 2021; Hayashi et al., 2021; Yokoyama et al., 2021).

In order to capitalize on the statistical benefits that simultaneous multi-slice (SMS) imaging has provided in human imaging (Feinberg et al., 2010; Feinberg and Setsompop, 2013; Risk et al., 2021), recent studies have begun exploring the use of SMS acquisitions with bespoke multichannel receive coils in imaging of anaesthetised NHPs (Autio et al., 2020). Yet, the differing biological conditions between awake and anaesthetised states could hinder utility of the results from the anaesthetised NHP experiments when compared to the awake human state. This has led to a surge in awake NHP studies recently (Seah et al., 2014).

However, accelerated fMRI in awake behaving NHPs poses some unique challenges. Although the animal’s head is typically mechanically stabilised in such experiments, motion in the behaving animal’s body, hands, jaw, and facial musculature causes strong, fluctuating B0 field inhomogeneities within the brain. These dynamic field inhomogeneities cause not only considerable image distortion but invalidate the correspondence between the imaging data and calibration data used for SMS image reconstruction. This results in additional ghosting and residual aliasing artifacts (from other simultaneously excited slices) that cannot be corrected in post processing and can degrade image quality.

Recently, we proposed a method to correct for these dynamic B0 changes (Shahdloo et al., 2022) by using the reference navigator data which are often available in most standard echo planar imaging (EPI) sequences used for fMRI data acquisition (Fig. 1a). This method uses navigator data acquired at every time-point for Nyquist ghost correction, and the GRAPPA-based operator weights trained on a calibration scan to estimate a first-order dynamic field perturbation at each time-point relative to the calibration scan. This can be interpreted as estimating a net translation in k-space due to the field perturbation. Then, this estimate is used to transform the acquired data to enhance consistency with the calibration data for a ghost- and alias-free SMS image reconstruction (Fig. 1b). While we have recently shown improvements in conventional image fidelity metrics, the impact of this off-resonance correction method has yet to be quantified on downstream estimates of brain activation.

**Figure 1.**
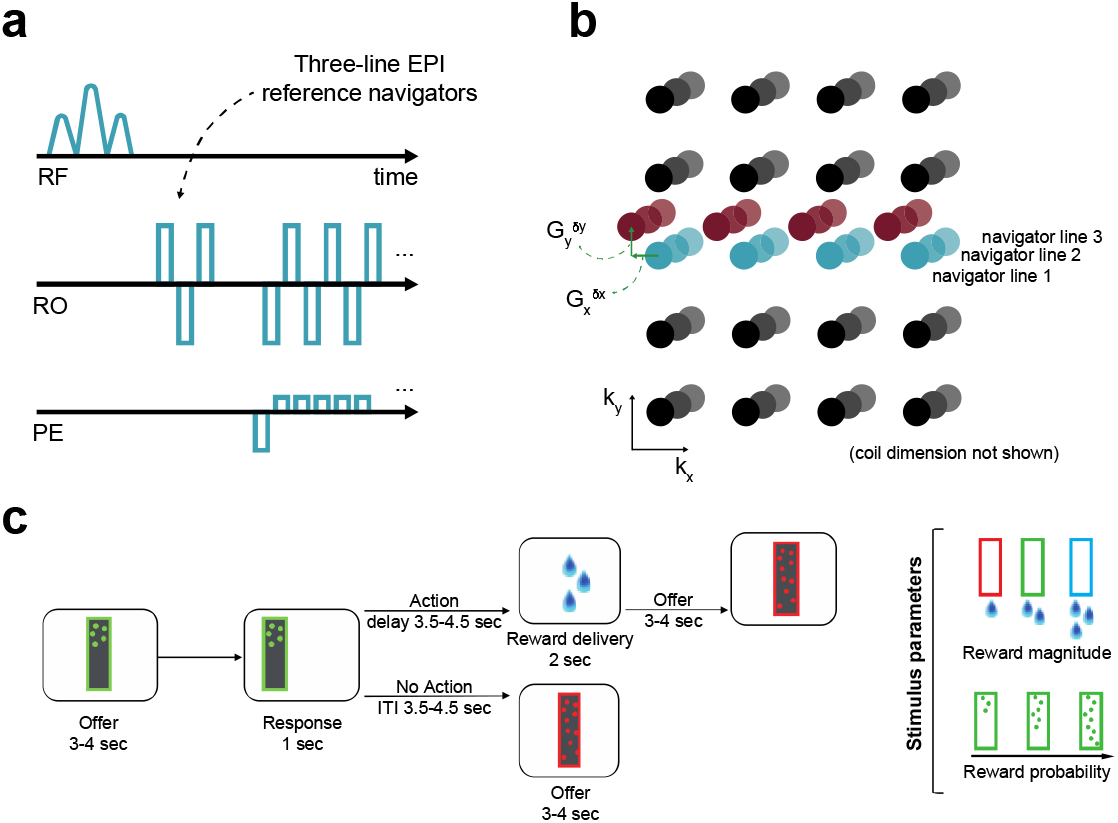
Off-resonance correction, and the experimental paradigm. **(a,b)** EPI reference navigators in each frame are compared to a reference frame to estimate the dynamic linear off-resonance. These estimates are then used to correct the imaging data (Shahdloo et al., 2022). **(c)** This off-resonance correction method was validated in a decision-making task in awake NHPs, where the animals decide to act based on the number and colour of dots appearing on the screen, respond by touching a pad, and receive a liquid reward based on the response. The experimental paradigm involves a wide range of body motion, as well as irregular events.

Here, we aim to validate this off-resonance correction method using GLM analysis of task-fMRI data from 40 awake behaving NHP scans across 4 animals. Our results show significant reductions in image bias and variance, as well as improvements in temporal signal to noise ratio (tSNR) and detected activations across the brain.

## Methods

### Behavioural task

In vivo 2D fMRI data from four male rhesus monkeys were collected (weight 14.1– 16.8 kg, 6-8 years of age) while they were performing a decision-making task. All procedures were conducted under licenses from the United Kingdom (UK) Home Office in accordance with The UK Animals Act 1986 (Scientific Procedures) and with the European Union guidelines (EU Directive 2010/63/EU).

The task was developed from the one originally used in our recent studies (Khalighinejad et al., 2022, 2020). At the beginning of each trial a frame (8 x 26 cm) appeared on the centre of the screen. The frame contained different number of dots. The colour and number of dots within the frame represented the magnitude and probability of potential reward, respectively. After 3-4s the frame moved to either the left or right side of the screen. Animals could respond (within 1s) by touching a custom-made infra-red touch sensor, on the side corresponding to the image. The probability of getting a reward and the drops of juice received depended on the number and colour of the dots, respectively (Fig. 1c). If they responded, they were offered drops of juice or no juice, based on reward probability on that trial. There was a delay of 3.5-4.5 s between response and outcome (action-outcome delay). If rewarded, drops of juice were delivered by a spout placed near the animal’s mouth. If they did not respond, the next offer appeared after a 3.5-4.5s intertrial interval (ITI). Reward magnitude and probability of the offer changed from one trial to another.

### Imaging protocol and reconstruction

The animals were head-fixed in sphinx position in an MRI-compatible chair in a clinical 3T scanner. Data were acquired using a 15-channel bespoke receive coil (Rapid Biomedical, Berlin, Germany), using the CMRR multi-band GRE-EPI sequence (Moeller et al., 2010; Setsompop et al., 2012) with parameters: TE/TR=25.4/1282ms, FA=63º, FOV=120mm, 1.25mm isotropic resolution, 42 slices, MB=2, in-plane acceleration factor R=2. Data from 10 runs were collected from each of the four animals, each containing 1213-1364 volumes. Raw data were separately reconstructed using the online reconstruction provided as part of the CMRR multi-band EPI sequence package and using our proposed GRAPPA-operator based dynamic off-resonance corrected reconstruction method. Single-band reference data and fully sampled calibration data were acquired along with the functional data as separate parts of the imaging sequence. Standard Nyquist ghost correction and dynamic zeroth-order B0 correction were applied on all reconstructions prior to off-resonance estimation and correction (Shahdloo et al., 2022).

### Preprocessing

Data preprocessing was performed using tools from FMRIB Software Library (FSL; Jenkinson et al., 2012), Advanced Normalization Tools (ANTs;18/10/2023 01:53:00 Tustison 2014), and the Magnetic Resonance Comparative Anatomy Toolbox (MrCat; Mars et al., 2016). A mean EPI reference image was created for each session, to which all volumes were non-linearly registered on a slice-by-slice basis along the phase-encoding direction to correct for time-varying distortions in the main magnetic field due to body and limb motion. The aligned and distortion-corrected functional images were then non-linearly registered to each animal’s high-resolution structural images. A group specific template was constructed by registering each animal’s structural image to the CARET macaque F99 space (Kolster et al., 2009). Finally, the functional images were temporally filtered (high-pass temporal filtering, 3-dB cutoff of 100s) and spatially smoothed (Gaussian spatial smoothing, full-width half maximum of 3mm). The same preprocessing parameters and pipelines were used for the uncorrected and off-resonance corrected reconstructions.

### Quantification and statistical analysis

#### Quantifying image quality

To validate the effect of dynamic off-resonance correction on image quality before preprocessing, the mean image difference compared to the single-band reference (bias) and the temporal coefficient of variation (variance) were calculated. To do this, the raw data were reconstructed using the vendor-provided reconstruction on the scanner, and with our proposed offline reconstruction that incorporates off-resonance correction. The single-band reference was reconstructed using the vendor-provided reconstruction on the scanner.

To assess the image quality improvement with finer granularity, tSNR values in anatomical regions of interest (ROIs) were compared between the uncorrected and corrected reconstructions. To enable pooling of tSNRs across animals and sessions, the data were mapped from the functional space onto the standard F99 space, where tSNR was taken as the ratio of the signal mean to standard deviation in each voxel. To create anatomical ROIs, voxel masks were created for each ROI in the F99 standard space using the Rhesus Monkey Brain Atlas (Paxinos, 2009). Masks were then transformed from the standard space to each participant’s structural space by applying a standard-to-structural warp field and from structural to functional space by applying a structural-to-functional affine matrix. Transformed masks were thresholded, binarised and were dilated by one voxel. tSNR values were averaged across voxels within anatomical ROI boundaries.

#### Whole-brain functional analysis

Whole-brain statistical analyses were performed at two-levels as implemented in FSL FEAT. At the first level, a univariate general linear model (GLM) was fit to the preprocessed data in each animal. The parameter estimates across sessions were then combined in a second-level mixed-effects analysis (FLAME 1+2), taking scanning sessions as random effect. The results were cluster-corrected with the voxel inclusion threshold of z=3.1 and cluster significance threshold of *p*=0.05. The data were pre-whitened to account for temporal autocorrelations.

The first-level analyses looked for voxels where the response reflected parametric variation in BOLD response according to the following GLM model

*BOLD* = β_0_ + β_1_ *offer* + β_2_ *action_onset* + β_3_ *motor_response* + β_4_ *outcome* where *BOLD* is a column vector containing the times-series data for a given voxel. *offer* is an unmodulated regressor representing the main effect of stimulus presentation (all event amplitudes set to one). *action_onset* is also an unmodulated regressor representing the go-cue (frame moving to either the left or right side of the screen). *motor_response* is a binary regressor representing response (pressing on the touch sensor) vs. no-response. *outcome* is an unmodulated regressor representing the main effect of outcome. Regressors were modelled as a boxcar function with constant duration of 0.1s convoluted with a hemodynamic response function (HRF) specific for monkey brains. Regressor 1 was time-locked to the onset of the trial (presentation of the offer). Regressors 2 and 3 were time-locked to the go-cue onset. Regressor 4 was time-locked to the onset of the reward outcome phase.

#### Reliability analysis

Reliability of the fMRI analyses using the uncorrected and corrected reconstructions were assessed and compared. In a 10-fold cross validation, experimental sessions were pooled across subjects and randomly divided into two groups. The second level analysis was then performed on each of the groups. Thresholded z-score values in anatomical ROIs were compared between the two groups and reliability was measured as the linear correlation coefficient between the z-score values. Furthermore, to ensure that the applied off-resonance correction has not introduced bias in the activation baseline, mean z-score values across ROI voxels were compared. Baseline bias was taken as the absolute difference between the mean z-score in each ROI between the two groups.

## Results

Figure 2a,b show reduced bias and temporal variance due to off-resonance correction, indicating increased reconstruction fidelity. Notably, presence of dynamic off-resonance had led to reconstruction artifacts in the original data that were not addressed using the conventional preprocessing pipeline that was applied. These artifacts are most prominent in the anterior and posterior regions, where half-FOV aliasing artifacts are most prominent. Figure 2c shows the histogram of the bias and variance values across brain voxels. Distribution of bias and variance in the corrected reconstruction is skewed towards zero, indicating larger number of voxels with low bias and variance compared to the uncorrected reconstruction.

**Figure 2.**
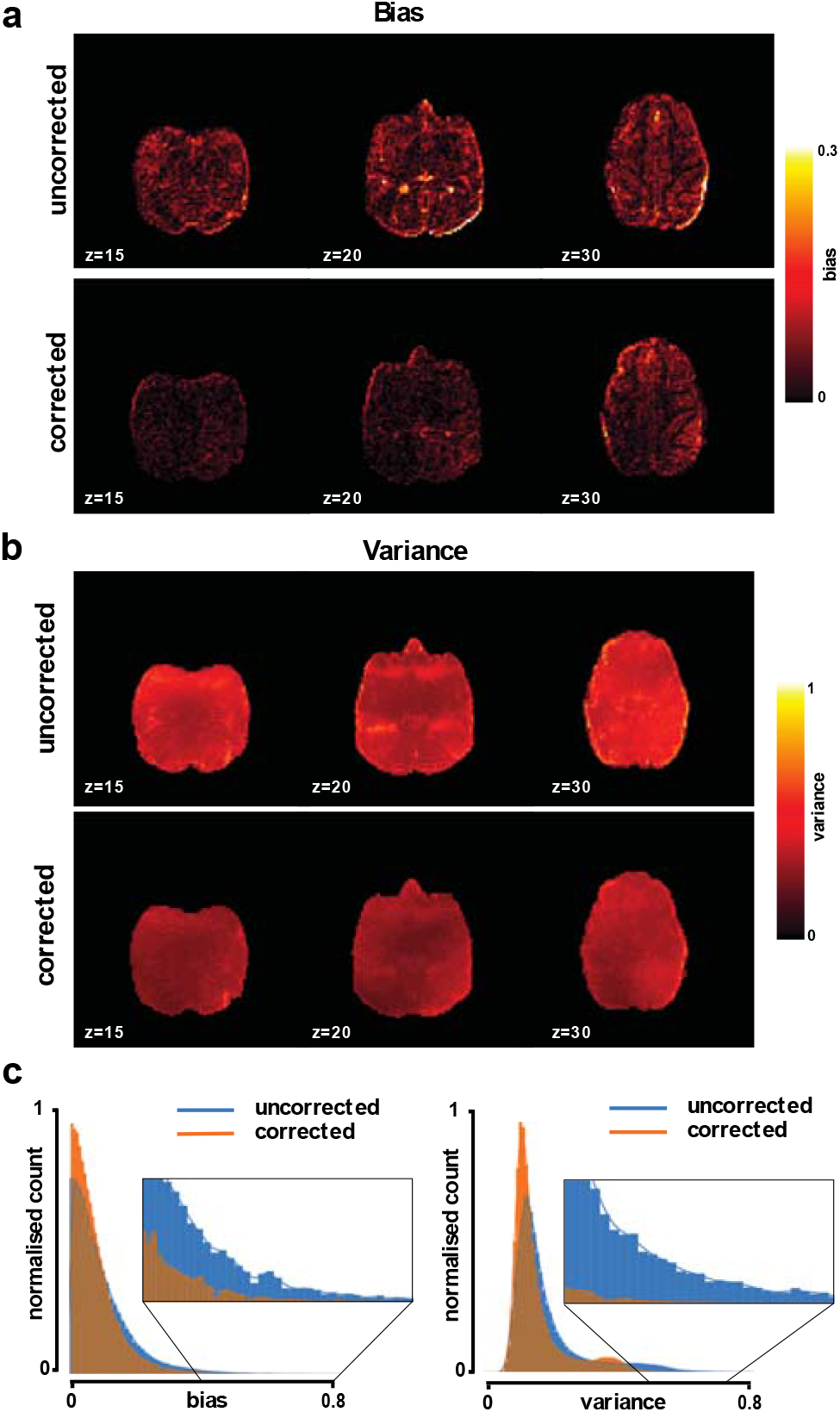
Reconstruction fidelity. Improvements in image quality were assessed using **(a)** reconstruction bias and **(b)** temporal variation, shown for a representative session. Off-resonance correction reduces the reconstruction bias and variance across the brain. **(c)** Histogram of these measures taken over brain voxels and pooled across sessions verifies the observed reduction in bias and variance.

Enhanced temporal signal stability would be expected as a result of the reduction in reconstruction variance. tSNR difference between corrected reconstruction compared to the uncorrected reconstruction was calculated and averaged in anatomical ROIs, shown in Fig. 3. tSNR is significantly increased in the studied ROIs (*q*(FDR)<0.05, non-significant in TCpol, PFCom, CCS).

**Figure 3.**
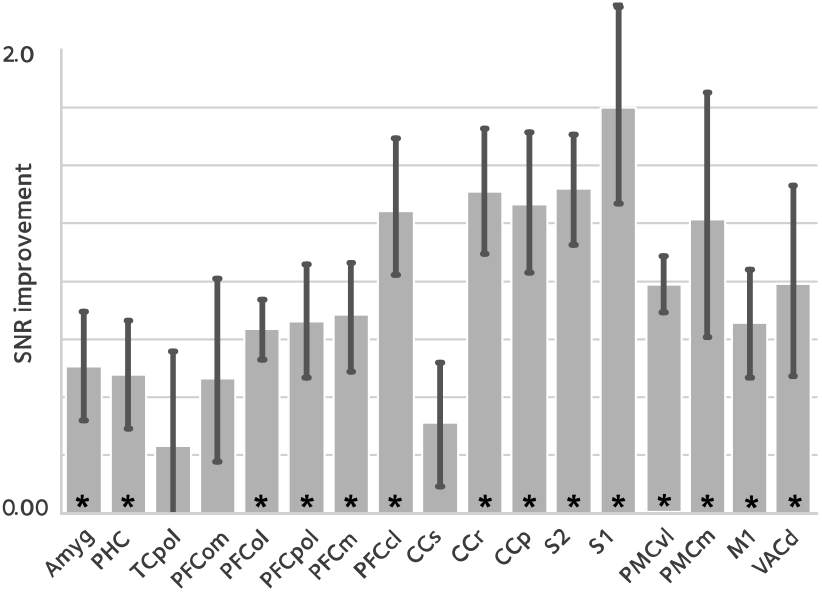
tSNR enhancement. To assess the effect of image quality improvements on the fMRI time series, temporal signal to noise ratio (tSNR) was compared between reconstructions, in anatomical ROIs (mean ± sem; asterisks denote statistical significance at q(FDR)<0.05). tSNR is significantly improved in most of the studied ROIs that cover the whole brain.

Here, we have hypothesised that the reduced bias and variance, and enhanced temporal signal stability should reflect the reduction in artifacts, and hence, increased statistical power in the downstream GLM analyses. To test this hypothesis, we performed first-level analysis. Figure 4 and 5 show thresholded z-score maps in representative sessions in two animals. Representative maps relating to the *offer* and *motor_response* regressors are shown, as they could elicit responses in widely differing brain regions. At a set threshold, the proposed off-resonance correction leads to a larger number of activated voxels, and larger effect sizes which is noticeable even at the level of single sessions.

**Figure 4.**
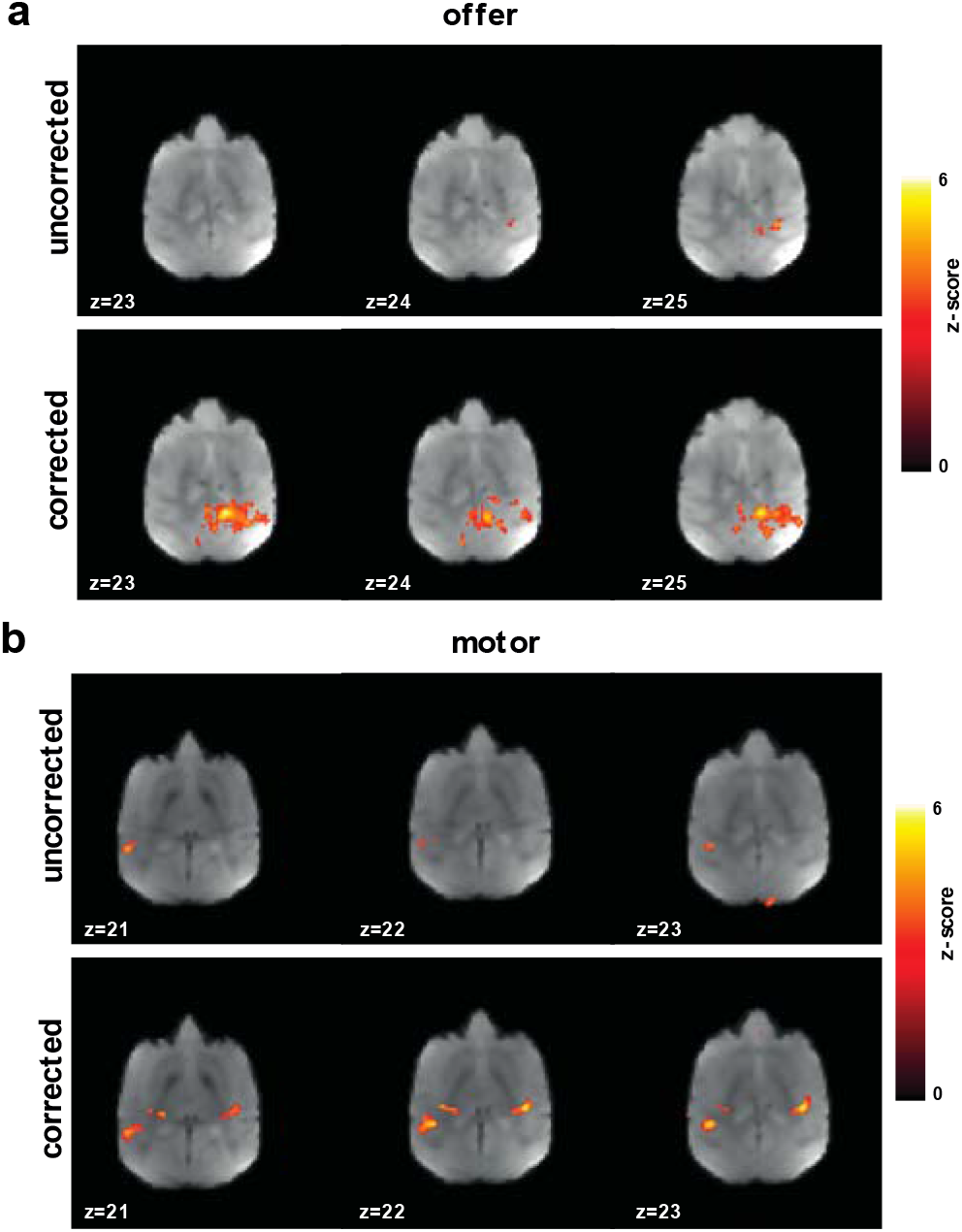
Improvement in first level GLM analysis. GLM models were fit to the data in each session to identify voxels that were activated **(a)** by *offer*, and **(b)** by the *motor_response* variables. Thresholded z-scores in a representative session are shown. Functional signal quality is improved using the off-resonance correction, as reflected by the larger number of voxels passing the threshold.

**Figure 5.**
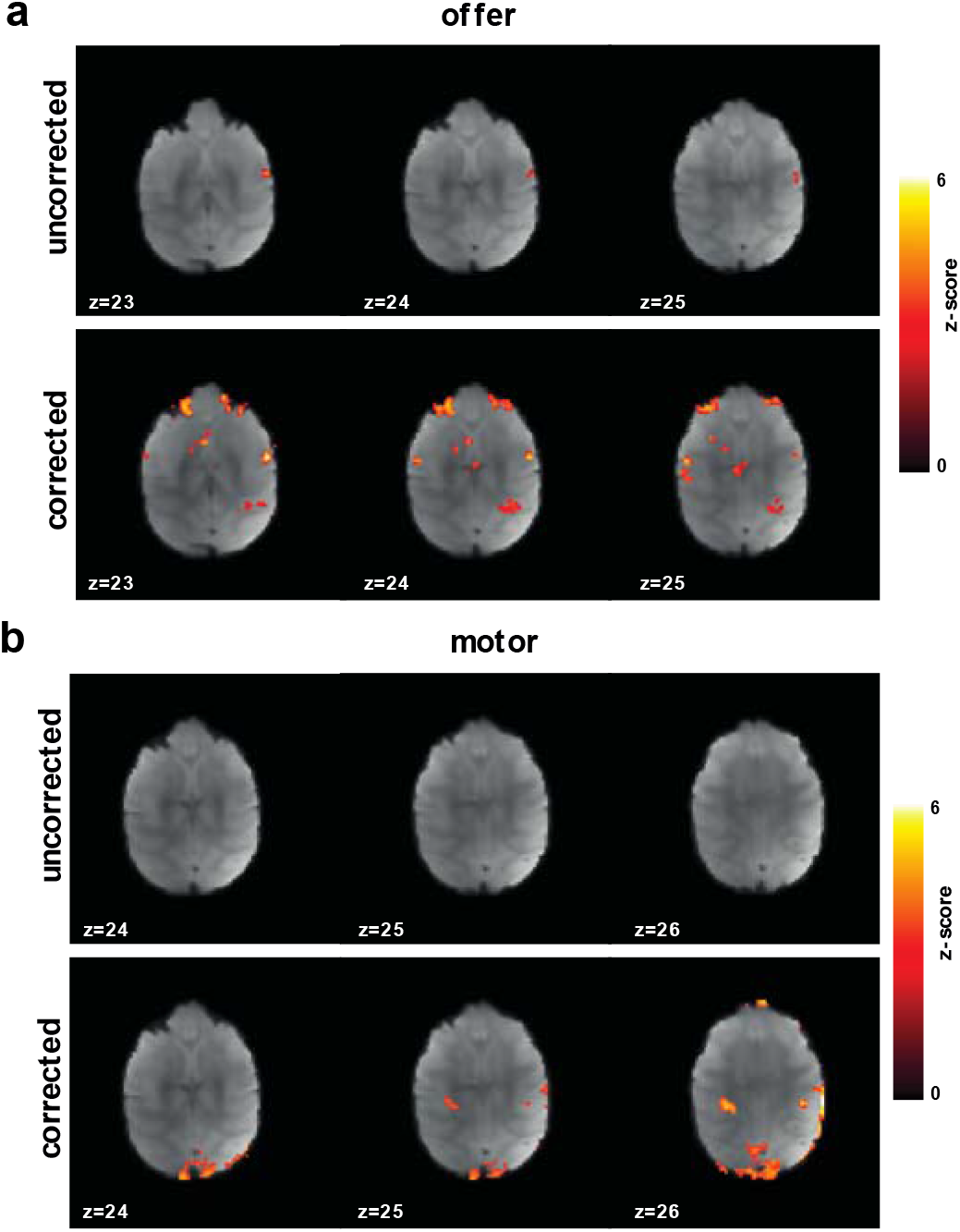
Improvement in first level GLM analysis. Thresholded z-score maps in a representative session from a second animal showing activated voxels **(a)** by *offer*, and **(b)** by the *motor_response* variables. The improvements in functional signal quality are consistent across animals.

Inherent noise at the single-session level could lead to high variation among sessions. Thus, we also investigated the results of a second-level mixed-effects analysis, taking scanning sessions as random effect. Figure 6 shows the 2^nd^-level z-score maps for the same regressors shown in Figs 4, 5 (thresholded at z=3.1). Reaffirming the results from the first-level analysis, off-resonance correction leads to a larger number of voxels surviving the threshold (Table I and II).

**Figure 6.**
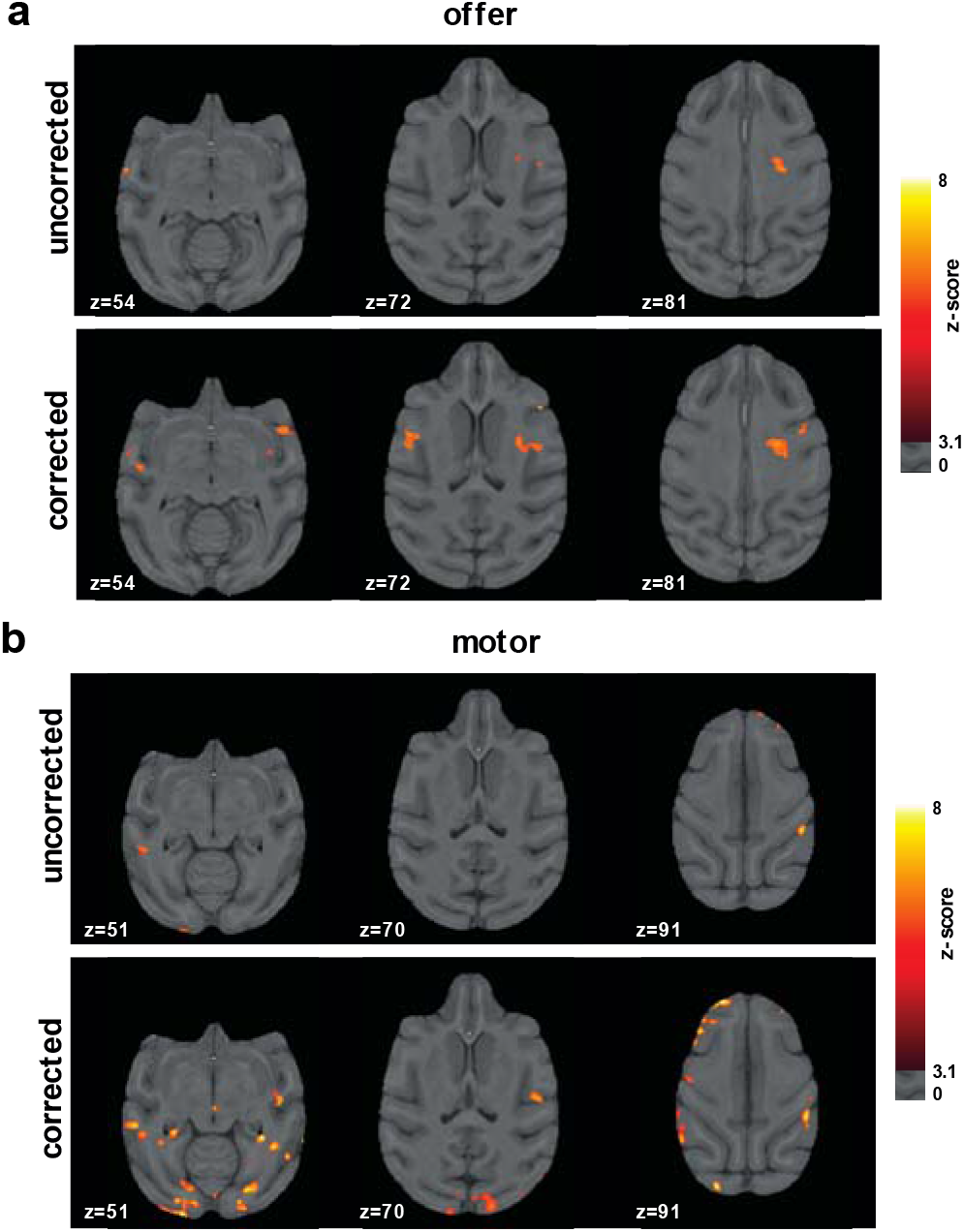
Improvements in second-level analysis. A second-level mixed-effects analysis was performed taking scanning sessions as random effect. Thresholded z-score maps are shown indicating activated voxels **(a)** by *offer*, and **(b)** by the *motor_response variables*. The off-resonance correction significantly improves the results of the higher-level GLM analysis.

**Table I.** Activated voxels by *offer*. To quantify the improvements in the higher-level analysis, the number of activated voxels by the *offer* variable were counted in each ROI. The off-resonance correction improves the number of activated voxels, as well as the peak z-scores across the brain.

|  | Uncorrected |  | Corrected |  |
| --- | --- | --- | --- | --- |
|  | N voxels | Max z | N voxels | Max z |
| *Amyg* | 130 | 3.77 | 1535 | 4.26 |
| *PHC* | 326 | 3.22 | 287 | 3.51 |
| *TCpol* | 0 | - | 416 | 3.43 |
| *CCr* | 0 | - | 215 | 3.28 |
| *CCp* | 209 | 3.12 | 701 | 4.07 |
| *S1* | 354 | 3.54 | 613 | 4.47 |
| *S2* | 107 | 3.57 | 887 | 4.14 |
| *PCm* | 20 | 3.30 | 69 | 3.62 |
| *PMCvl* | 444 | 3.19 | 3947 | 4.92 |
| *PMCm* | 0 | - | 1127 | 3.92 |
| *VACd* | 105 | 3.11 | 374 | 3.38 |

**Table I.** Activated voxels by the *motor_response*. To quantify the improvements in the higher-level analysis, the number of activated voxels by *motor_response* variable were counted in each ROI. Consistent with Table I, the off-resonance correction improves the number of activated voxels, as well as the peak z-scores across the brain.

|  | Uncorrected |  | Corrected |  |
| --- | --- | --- | --- | --- |
|  | N voxels | Max z | N voxels | Max z |
| *Amyg* | 0 | - | 862 | 3.80 |
| *TCpol* | 0 | - | 640 | 3.85 |
| *CCp* | 160 | 3.78 | 191 | 4.58 |
| *S1* | 178 | 3.32 | 479 | 4.56 |
| *S2* | 80 | 3.63 | 907 | 3.69 |
| *PMCvl* | 3753 | 5.76 | 555 | 3.68 |
| *PMCm* | 0 | - | 297 | 3.92 |
| *M1* | 0 | - | 254 | 3.78 |

The experimental procedure used here results in sessions with varying time lengths. Thus, we investigated the reliability of the second-level analysis by correlating the second-level z-score maps between two random splits of the sessions pooled across subjects, via a cross-validation procedure. Figure 7a shows the second-level reliability in anatomical ROIs. Off-resonance corrected reconstruction leads to more reliable results in ROIs with activated voxels.

**Figure 7.**
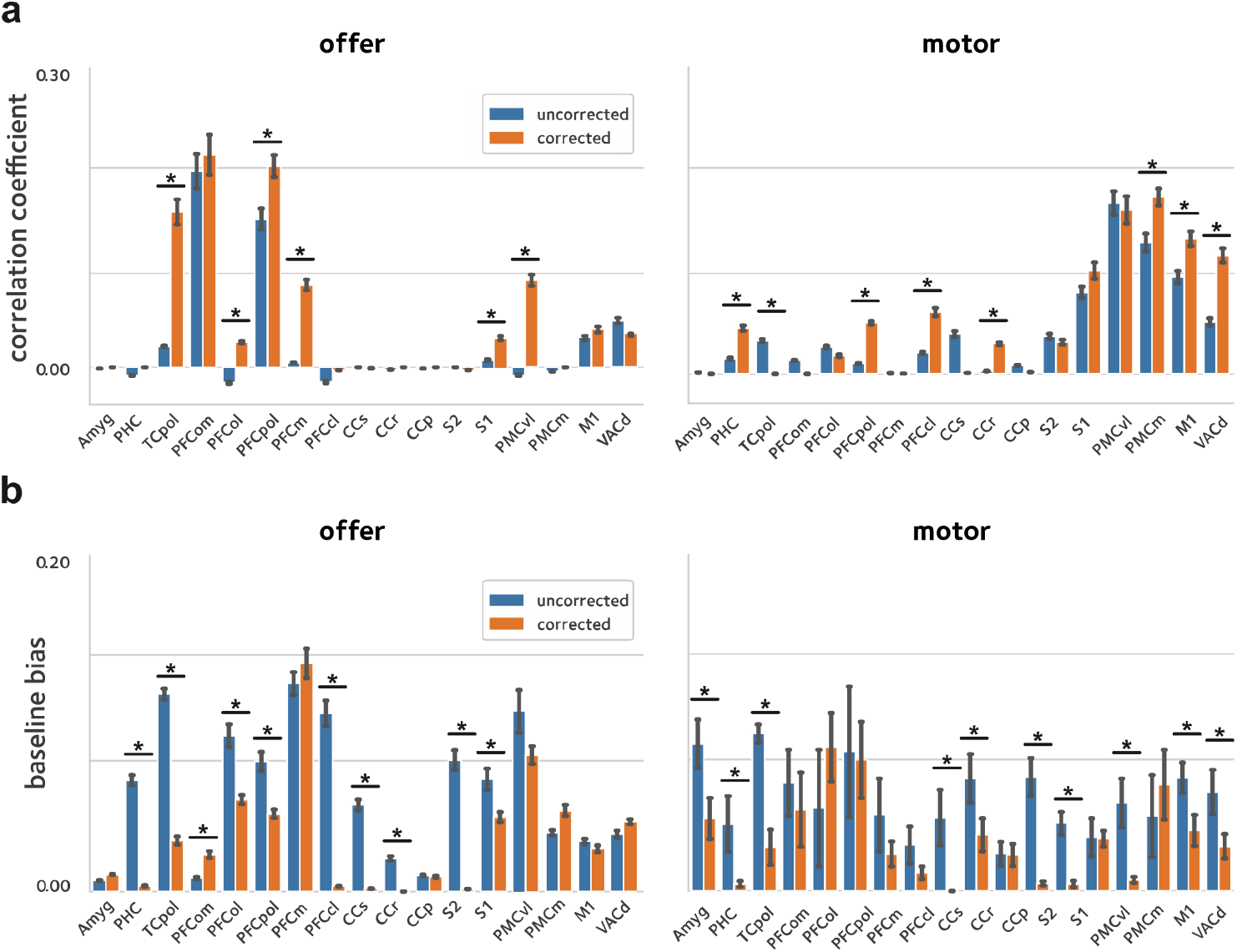
Reliability of activation estimates. To quantify the improvements in reliability of the estimated activations, a cross-validation procedure was used were the sessions pooled across animals were randomly divided in two and models were fit in each sub-group separately. Reliability was quantified by taking the **(a)** correlation, and **(b)** baseline bias between the estimates in the two sub-groups (asterisks denote statistical significance at q(FDR)<0.05). The off-resonance correction significantly improves the reliability of activation estimates across the brain.

## Discussion

Accelerated awake NHP fMRI entails unique challenges compared to human fMRI, one of the main ones being reconstruction artifacts due to dynamic off-resonance changes. In head-posted animals, motion in body parts could cause strong field changes that introduce artifacts and distort the images in a way that cannot be corrected in preprocessing. There is therefore a need for approaches for artifact suppression and distortion correction during image reconstruction. Recently, we proposed a method to achieve this using the navigator data that is already collected during the fMRI acquisition. Here, we used a large amount of awake NHP data from a physically demanding and dynamic decision-making experiment to investigate the efficacy of this off-resonance correction. Our results show that improvements in image fidelity and temporal stability lead to improved estimates of brain activation. Moreover, the method studied here relies only on the navigator data acquired using conventional accelerated acquisition sequences. Thus, applying this off-resonance correction retrospectively to previously acquired scans is possible and can improve fMRI analyses if raw imaging data are available.

The off-resonance correction method used here assumes first-order spatial off-resonance perturbations that have been shown to be a good approximation in imaging headposted NHPs (Pfeuffer et al., 2007). One limitation of this correction is that the accuracy of the first-order approximation can potentially suffer from extreme body motion. The experimental paradigm used here entails long runs where animals perform choices by pressing on a touch sensor and receive liquid rewards, each of which involve a large range of motion in body parts. The results show that the linear correction can significantly improve parameter estimates across the brain. However, highly dynamic experimental setups may benefit from extra nonlinear preprocessing steps to reduce the residual artifacts and distortion.

It should also be noted that the correction performance depends heavily on the capability of the receive coil in parallel imaging. The data used here were acquired using a bespoke NHP coil with relatively good spatial coverage. However, the improvements might be hindered if an unsuitable coil had been used for data collection.

In conclusion, the method validated here enables more robust and reliable NHP imaging in accelerated acquisitions that leads to significantly improved functional analysis results in awake NHP studies.

## Acknowledgements

The Wellcome Centre for Integrative Neuroimaging is supported by core funding from the Wellcome Trust (203139/Z/16/Z). NK is funded by BBSRC Discovery Fellowship (BB/W008947/1). MR is supported by BBSRC fellowship (BB/W003392/1) and Wellcome Trust award (221794/Z/20/Z). MC is supported by the Canada Research Chairs Program.

## Notes

### Competing Interest Statement

The authors have declared no competing interest.

